# The TWEAK/Fn14 signaling promotes skeletal muscle wasting during cancer cachexia

**DOI:** 10.1101/2024.10.06.616866

**Authors:** Meiricris Tomaz da Silva, Anirban Roy, Anh Tuan Vuong, Aniket S. Joshi, Cristeena Josphien, Meghana V Trivedi, Sajedah M Hindi, Vihang Narkar, Ashok Kumar

## Abstract

Cachexia is an involuntary loss of body weight mostly due to skeletal muscle wasting. The proinflammatory cytokine TWEAK and its receptor Fn14 constitute a major signaling system that regulates skeletal muscle mass in diverse conditions. However, the role of TWEAK/Fn14 system in the regulation of skeletal muscle mass during cancer-induced cachexia remains poorly understood. In this study, we demonstrate that the levels of Fn14, but not TWEAK, are induced in skeletal muscle of multiple mouse models of cancer cachexia. Targeted deletion of Fn14 inhibits muscle wasting and gene expression of multiple components of the ER stress-induced unfolded protein response (UPR) in the KPC mouse model of pancreatic ductal adenocarcinoma (PDAC) cancer cachexia. The TWEAK/Fn14 signaling activates PERK and IRE1α arm of the UPR and inhibits protein synthesis in cultured primary myotubes. Inhibition of PERK using pharmacological or molecular approaches improves protein synthesis and inhibits atrophy in TWEAK-treated cultured myotubes. Silencing of Fn14 in KPC cells prior to their inoculation in pancreas of mice also attenuates tumor growth without having any significant effect on muscle atrophy. The knockdown of Fn14 inhibits proliferation, migration, and invasion of cultured KPC cells. Finally, our results demonstrate that targeted ablation of Fn14 also attenuates muscle atrophy in the Lewis lung carcinoma model of cancer cachexia. Altogether, our study provides initial evidence that the inhibition of TWEAK/Fn14 signaling can prevent tumor growth and skeletal muscle wasting during cancer-induced cachexia.

## INTRODUCTION

Cancer cachexia is a debilitating multifaceted syndrome characterized by the involuntary loss of body weight and skeletal muscle mass, with or without a concurrent loss of adipose mass (1). About two-thirds of patients with advanced cancer develop cachexia, which drastically reduces their quality of life, increases resistance to immunotherapy and chemotherapy, and decreases the overall survival rates (2, 3). However, at present there are no effective drugs or therapeutic approaches to reverse the condition in cancer patients. Although the mechanisms underlying cancer cachexia remain poorly understood, it has been consistently observed that various factors released from the tumor itself and/or the host in response to the tumor growth contribute to the loss of muscle mass in both preclinical animal models and cancer patients (2, 4–6). Accumulating evidence also suggests that inhibiting muscle wasting not only improves survival but also slows down tumor growth by depriving the tumor of key energy substrates essential for its rapid proliferation (7).

Skeletal muscle wasting in many conditions, including cancer cachexia occurs due to an imbalance between the rates of muscle protein synthesis and degradation. The ubiquitin-proteasome system is one of the most important proteolytic systems, which is responsible for the degradation of bulk of muscle proteins during cancer-induced cachexia (8, 9). Accumulating evidence also suggests that endoplasmic reticulum (ER) stress-induced unfolded protein response (UPR) plays an important role in the regulation of skeletal muscle mass in various conditions (10, 11). The UPR is controlled by three ER transmembrane sensors: protein kinase R-like endoplasmic reticulum kinase (PERK), inositol-requiring protein 1α (IRE1α), and activating transcription factor 6 (ATF6). In response to ER stress, PERK is activated, which then phosphorylates eukaryotic initiation factor-2α (eIF2α) protein, resulting in general inhibition of protein synthesis while augmenting the translation of selective molecules, such as ATF4. Similarly, stress in ER also leads to the activation of IRE1α endonuclease activity which induces the alternative splicing of X-box protein 1 (XBP1) mRNA leading to the formation of a transcriptionally active spliced XBP1 (sXBP1) protein. The sXBP1 transcription factor induces the gene expression of various chaperones and other proteins, which ultimately restore proteostasis in the ER (12, 13). While the activation of the UPR is a physiological response to improve protein folding capacity within the ER, unmitigated and sustained activation of the UPR can lead to deleterious consequences, such as inflammation, insulin resistance, and cell death (12, 13). Recent studies have also demonstrated that several components of the UPR are induced in skeletal muscle of animal models of cancer cachexia (14, 15). However, the mechanisms underlying the UPR activation and its potential role in the regulation of skeletal muscle mass during cancer cachexia remain less understood.

Systemic inflammation plays an important role in skeletal muscle wasting in many chronic disease states, including cancer (2, 3, 8). TNF-like weak inducer of apoptosis (TWEAK, gene name: *Tnfsf12*) is a proinflammatory cytokine belonging to TNF superfamily. TWEAK functions through binding to fibroblast growth factor-inducible 14 (Fn14, gene name: *Tnfrsf12a*), a member of the TNF receptor superfamily. TWEAK, which is constitutively expressed in many cell types, is initially synthesized as a type II transmembrane protein, but it can be cleaved by furin to produce a soluble cytokine (16–18). In contrast, the Fn14 receptor is expressed at relatively low levels in healthy tissues. The activity of the TWEAK-Fn14 axis is enhanced due to a highly induced local expression of Fn14 in injured tissues and many disorders including autoimmune diseases, cancers, and neuromuscular disorders, triggering activation of multiple downstream signaling pathways to remodel these tissues (16). Increased expression of Fn14 has been detected in many types of cancers, including glioma, melanoma, and lung and breast cancers to promote cell proliferation, migration, invasion, and tumor metastasis (19–22). Furthermore, overexpression of Fn14 is associated with a worse clinical outcome in advanced stage of multiple cancers (23, 24).

The TWEAK/Fn14 system has now emerged as an important mediator of skeletal muscle wasting in diverse conditions (25). Transgenic overexpression of TWEAK induces skeletal muscle wasting and metabolic abnormalities in mice (26, 27). Intriguingly, the levels of Fn14, but not TWEAK itself, are induced in skeletal muscle in multiple catabolic conditions, including functional denervation, fasting, burn injury, spinal cord injury-associated muscle atrophy, and many other degenerative muscle disorders (25, 28–31). We previously reported that genetic ablation of TWEAK inhibits muscle wasting in response to denervation (28). Moreover, muscle-specific deletion of Fn14 improves exercise capacity and prevents denervation-induced muscle atrophy in mice (32). Interestingly, using antibody-based approaches to neutralize Fn14, a previously published study showed that Fn14 signaling in tumor cells is responsible for muscle wasting in an animal model of cancer cachexia (33). However, the cell-autonomous role of Fn14 in skeletal muscle wasting in tumor-bearing hosts remains completely unknown.

In this study, using muscle-specific Fn14 knockout mice, we investigated the role of Fn14 signaling in the regulation of skeletal muscle mass and growth of KRAS^G12D^ P53^R172H^ Pdx Cre^+/+^ (KPC) tumor, a mouse model of pancreatic ductal adenocarcinoma (PDAC) cancer cachexia. Our results demonstrate that myofiber-specific deletion of Fn14 inhibits the loss of skeletal muscle mass in KPC tumor-bearing mice. Fn14 signaling mediates the growth of KPC tumor *in vivo* and augments the proliferation, migration, and invasiveness of cultured KPC cells. Furthermore, our results suggest that the TWEAK/Fn14 signaling activates ER stress/UPR in skeletal muscle during cancer cachexia and the activation UPR is an important mechanism for the inhibition of protein synthesis and induction of atrophy in TWEAK-treated cultured myotubes.

## RESULTS

### Fn14 levels are increased in models of cancer cachexia

We first examined mRNA levels of TWEAK and Fn14 in skeletal muscle of KPC model of pancreatic cancer cachexia (34). Results showed that the mRNA levels of Fn14, but not TWEAK, were significantly increased in skeletal muscle of KPC tumor-bearing mice compared to corresponding control mice (**Fig. 1A**). We next measured mRNA levels of TWEAK and Fn14 in 2-month old Apc^Min/+^ mice, a model of colorectal cancer cachexia (35). Similar to KPC model, the mRNA levels of Fn14 were also significantly elevated in skeletal muscle of Apc^Min/+^ mice compared to corresponding wild-type mice. By contrast, mRNA levels of TWEAK remained comparable between wild type and Apc^Min/+^ mice (**Fig. 1B**). Western blot analysis showed that protein levels of Fn14 were also increased in skeletal muscle of Apc^Min/+^ mice compared to wild-type mice (**Fig. 1C**). We also analyzed a publicly available RNA-Seq dataset (GSE142455) studying global gene expression in skeletal muscle of C26 colorectal cancer mouse model (36). There was a significant increase in the mRNA levels of Fn14 in skeletal muscle of mice injected with C26 tumor cells subcutaneously or intrasplenically compared to corresponding control mice (**Fig. 1D**). Altogether, these results suggest that gene expression of Fn14 is induced in skeletal muscle of multiple models of cancer cachexia.

**FIGURE 1.**
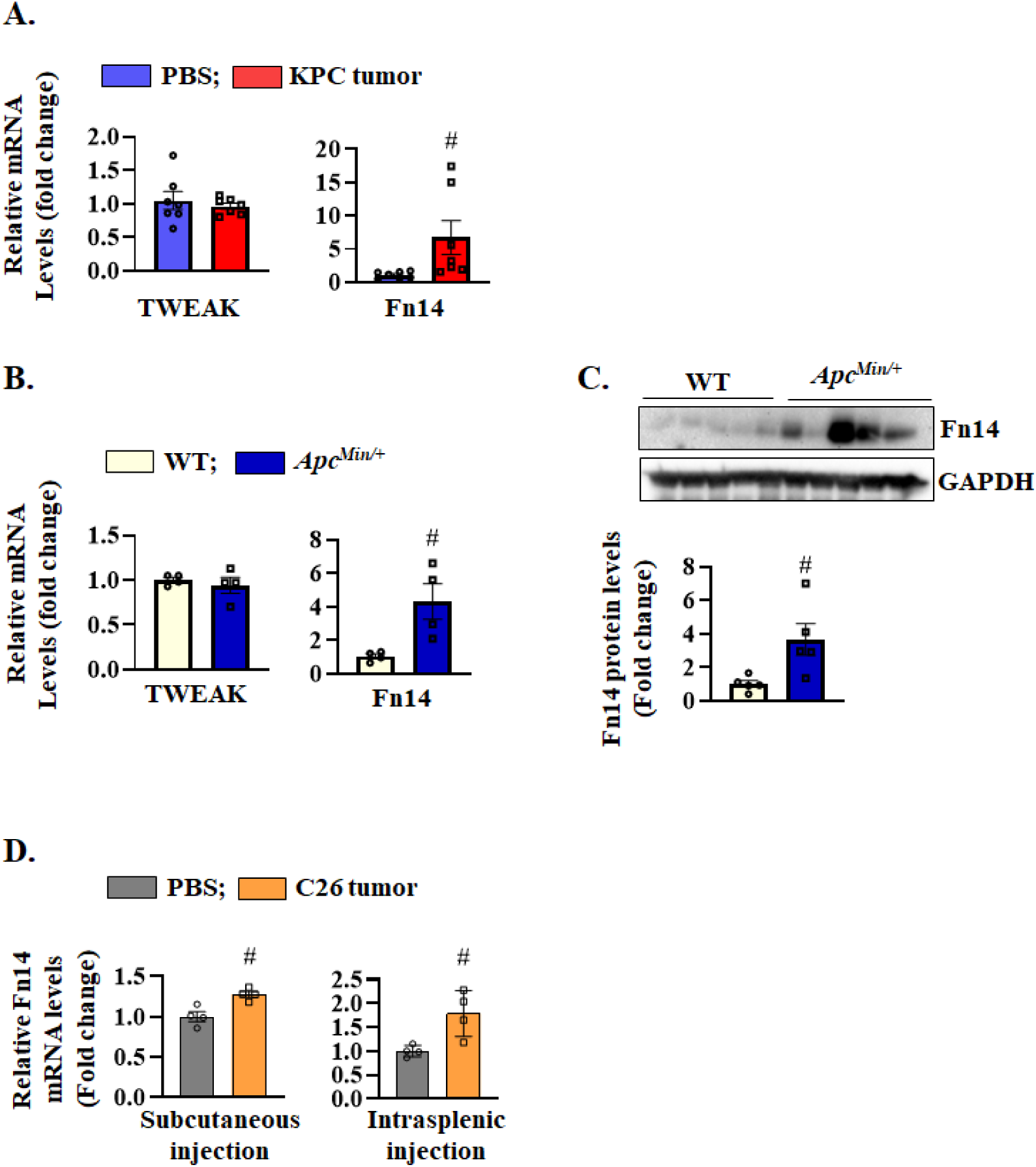
Levels of Fn14 in skeletal muscle of mouse models of cancer cachexia. **(A)** Relative mRNA levels of TWEAK and Fn14 in quadriceps muscle of control and KPC tumor-bearing mice. n = 7 mice in each group. **(B)** Relative mRNA levels of TWEAK and Fn14 in gastrocnemius muscle of 2-month old WT and Apc^Min/+^ mice. **(C)** Immunoblot and densitometry analysis of Fn14 protein levels in GA muscle of WT and Apc^Min/+^ mice. n = 5 mice in each group. **(D)** Relative Fn14 mRNA levels in quadriceps muscle of control (injected PBS alone) and C26 tumor-bearing (injected subcutaneously or intrasplenically) mice analyzed from a publicly available bulk RNA-Seq dataset (GSE142455). n = 4 mice in each group. All data are presented as mean ± SEM and analyzed by unpaired Student *t* test. ^#^p ≤ 0.05, values significantly different from WT mice or corresponding control mice injected with PBS alone.

### Target deletion of Fn14 inhibits muscle wasting in KPC tumor-bearing mice

To understand the role of Fn14 in the regulation of skeletal muscle mass and function during cancer cachexia, we crossed floxed Fn14 (i.e., Fn14^fl/fl^) mice with MCK-Cre line to generate muscle-specific Fn14 knockout (i.e., Fn14^mKO^) and control (i.e., Fn14^fl/fl^) mice as described (32). Next, 10-week old littermate male Fn14^fl/fl^ and Fn14^mKO^ mice were injected with phosphate buffer saline (PBS, served as control) or KPC cells in the pancreas. On day 26 after implantation of KPC cells, the mice were analyzed for body weight and grip strength. A significant reduction in tumor-free body weight and four-paw grip strength was observed in Fn14^fl/fl^, but not Fn14^mKO^ mice, in response to KPC tumor growth compared to corresponding control mice (**Fig. 2A, B**). Indeed, grip strength of KPC tumor-bearing Fn14^mKO^ mice was significantly higher compared to KPC tumor-bearing Fn14^fl/fl^ mice (**Fig. 2B**). Moreover, the wet weight of individual tibialis anterior (TA), soleus (Sol), and gastrocnemius (GA) muscle was also higher in KPC tumor-bearing Fn14^mKO^ mice compared to corresponding Fn14^fl/fl^ mice (**Fig. 2C**). By contrast, there was no significant difference in wet tumor weight between Fn14^fl/fl^ and Fn14^mKO^ mice (**Fig. 2D**). We next generated transverse sections of TA muscle and performed H&E staining and anti-laminin (to mark myofiber boundary) and DAPI staining followed by quantitative analysis of myofiber size (**Fig. 2E, F**). We found that the proportion of myofibers with larger cross-sectional area (CSA) was considerably enhanced in TA muscle of KPC tumor-bearing Fn14^mKO^ mice compared to Fn14^fl/fl^ mice (**Fig. 2G**). Indeed, average myofiber CSA in TA muscle was significantly higher in KPC tumor-bearing Fn14^mKO^ mice compared to KPC tumor-bearing Fn14^fl/fl^ mice (**Fig. 2H**).

**FIGURE 2.**
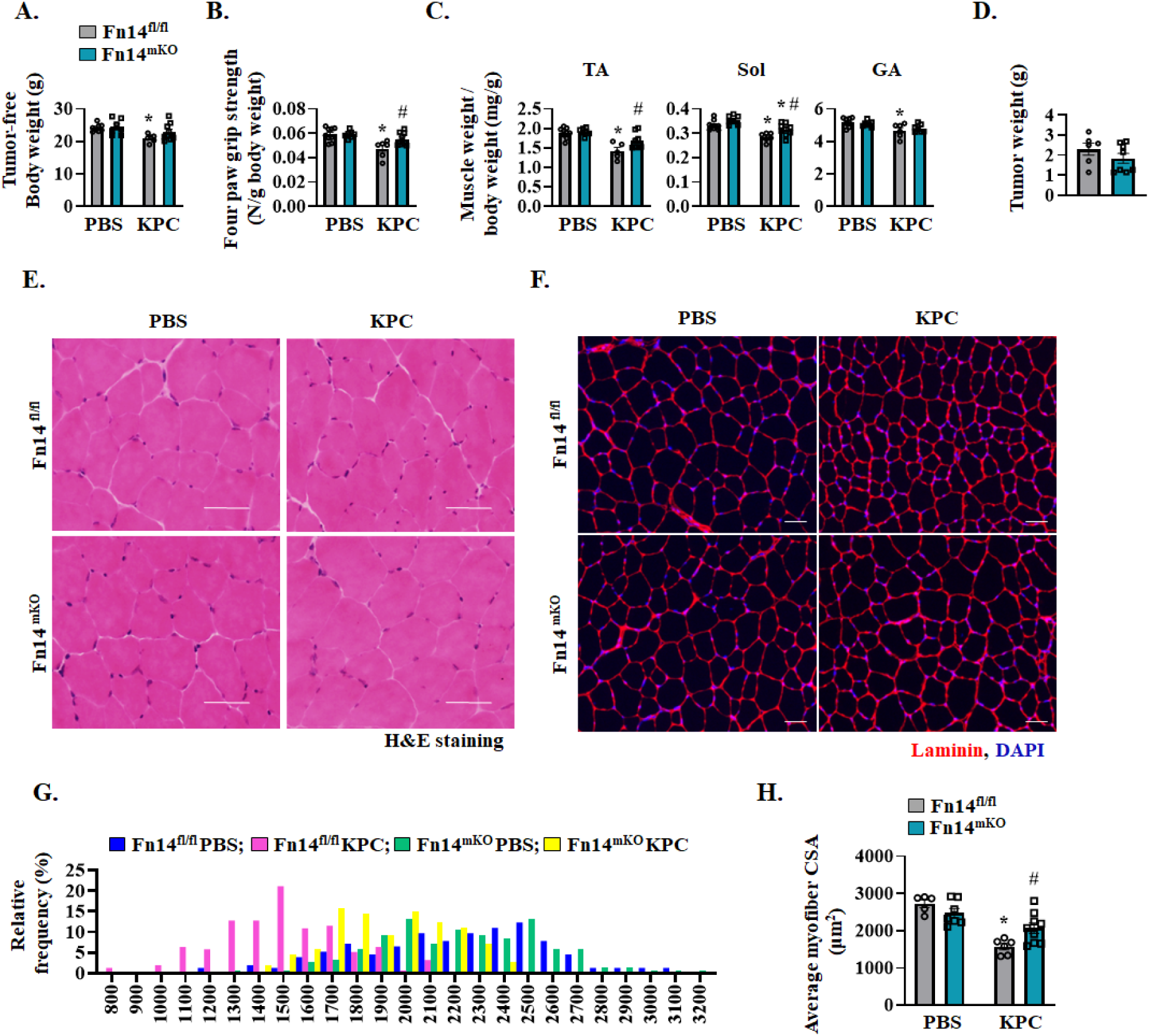
Genetic ablation of Fn14 inhibits TA muscle wasting in KPC tumor-bearing mice. 10-week old male Fn14^fl/fl^ and Fn14^mKO^ mice were injected with KPC cells or PBS alone (control) in the pancreas and monitored for 26 days. **(A)** Tumor-free body weight of control and KPC-tumor bearing Fn14^fl/fl^ and Fn14^mKO^ mice. **(B)** Quantification of average four-paw grip strength of control and KPC tumor-bearing Fn14^fl/fl^ and Fn14^mKO^ mice normalized by body weight. **(C)** Wet weight of tibialis anterior (TA), soleus (Sol), and gastrocnemius (GA) muscle normalized with tumor-free body weight. **(D)** Wet tumor weight. Representative photomicrographs of **(E)** H&E-stained, and **(F)** anti-laminin and DAPI-stained TA muscle sections of control and KPC tumor-bearing Fn14^fl/fl^ and Fn14^mKO^ mice. Scale bars, 50 μm. **(G)** Frequency distribution of myofiber cross-section area (CSA) in control and KPC tumor-bearing Fn14^fl/fl^ and Fn14^mKO^ mice. **(H)** Average myofiber CSA in TA muscle of control and KPC tumor-bearing Fn14^fl/fl^ and Fn14^mKO^ mice. n = 5-9 in each group. All data are presented as mean ± SEM and analyzed by two-way ANOVA followed by Tukey’s multiple comparison test or unpaired Student *t* test. *p ≤ 0.05, values significantly different from corresponding control mice. ^#^p ≤ 0.05, values significantly different from KPC-injected Fn14^fl/fl^ mice.

TA muscle of mice contains predominantly fast-type myofibers whereas soleus muscle contains both slow and fast-type myofibers. To understand whether the effect of deletion of Fn14 on myofiber size during cancer cachexia depends on muscle fiber type, we also generated muscle sections of soleus muscle followed by performing H&E staining and anti-laminin and DAPI staining (**Fig. 3A, B**). Quantitative analysis showed that the average myofiber CSA of soleus muscle of KPC tumor-bearing Fn14^mKO^ mice was also significantly higher compared to corresponding KPC tumor-bearing Fn14^fl/fl^ mice (**Fig. 3C, D**).

**FIGURE 3.**
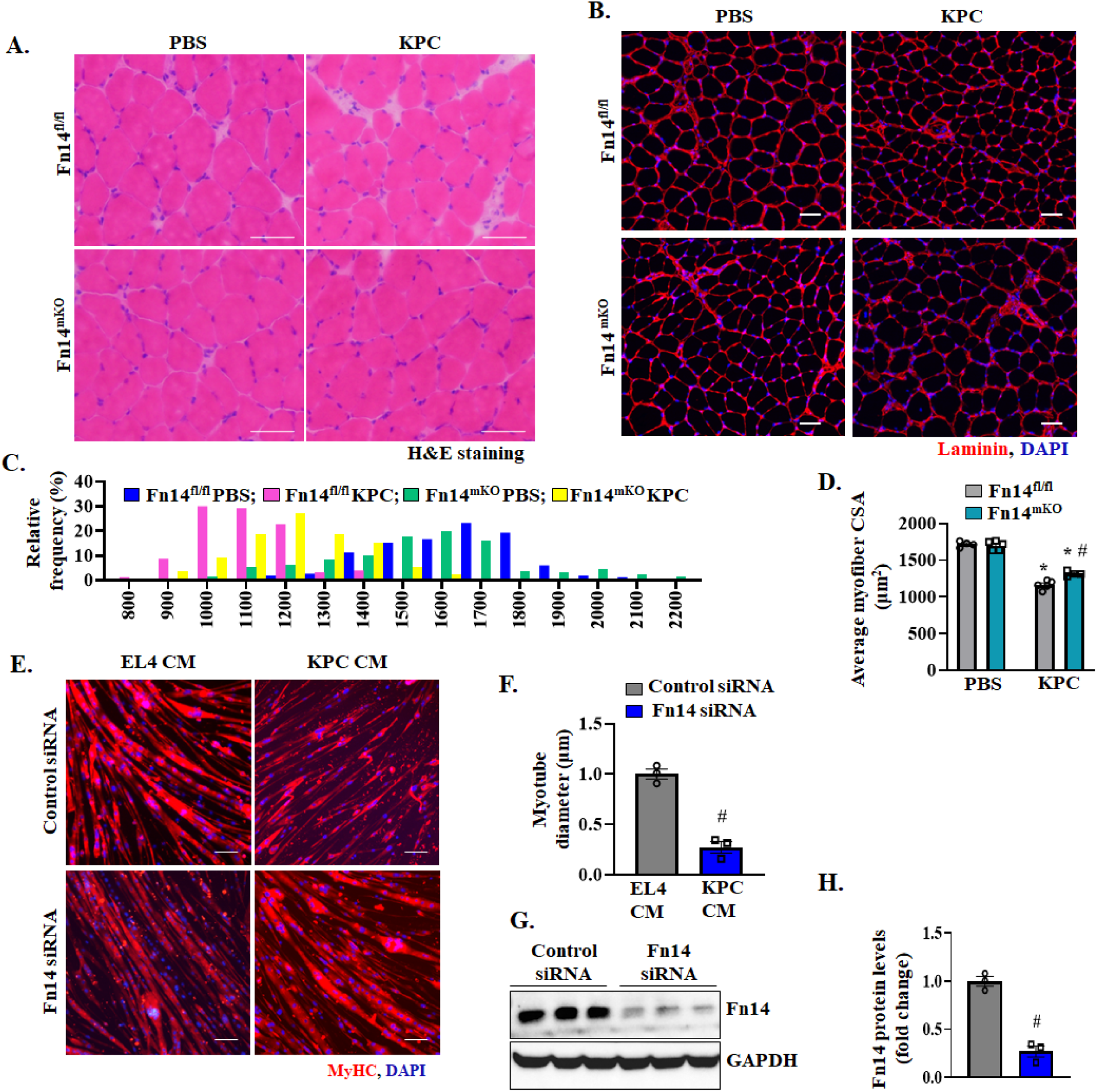
Inhibition of Fn14 attenuates atrophy in soleus muscle of mice and cultured myotubes. 10-week old male Fn14^fl/fl^ and Fn14^mKO^ mice were inoculated with KPC cells in the pancreas and monitored for 26 days. **(A)** Representative H&E-stained images of soleus muscle sections of control and KPC tumor-bearing Fn14^fl/fl^ and Fn14^mKO^ mice. Scale bars, 50 μm. **(B)** Representative anti-laminin-stained images of soleus muscle sections of control and KPC tumor-bearing Fn14^fl/fl^ and Fn14^mKO^ mice. Scale bars, 100 μm. **(C)** Relative frequency distribution of myofiber CSA in PBS and KPC tumor-bearing Fn14^fl/fl^ and Fn14^mKO^ mice. **(D)** Average myofiber CSA in control and KPC tumor-bearing Fn14^fl/fl^ and Fn14^mKO^ mice. n = 4-5 mice in each group. All data are presented as mean ± SEM and analyzed by two-way ANOVA followed by Tukey’s multiple comparison test. *p ≤ 0.05, values significantly different from skeletal muscle of corresponding control mice. ^#^p ≤ 0.05, values significantly different from KPC tumor-bearing Fn14^fl/fl^ mice. **(E)** WT myotubes were transfected with control or Fn14 siRNA for 24 h. The cells were washed and incubated with EL4 (non-cachectic) or KPC cells conditioned media (CM) at a 1:4 ratio for an additional 48 h. Representative images of myotube cultures after immunostaining for MyHC. DAPI was used to stain nuclei. Scale bars, 50 μm. **(F)** Quantification of average myotube diameters in control and Fn14 knocked down myotube cultures incubated with CM from EL4 or KPC cells. n = 3 biological replicates in each group. *p ≤ 0.05, values significantly different from corresponding EL4 CM treated myotubes. #p ≤ 0.05, values significantly different from KPC CM treated myotubes transfected with control siRNA. **(G)** Representative immunoblot and **(H)** densitometry analysis of Fn14 protein in myotube cultures transfected with control or Fn14 siRNA. n = 3 biological replicates in each group. #p ≤ 0.05, values significantly different from myotubes transfected with control siRNA.

We also determined the role of Fn14 in KPC tumor-induced muscle atrophy using cultured myotubes. We first prepared conditioned media (CM) from KPC cells and non-cachectic EL4 (a mouse T-cell lymphoma cell line) cells. Mouse primary myotubes were transfected with control or Fn14 siRNA followed by treatment with CM prepared from EL4 or KPC cells for 24 h. The cultures were then fixed and immunostained for MyHC protein. As expected, treatment with CM of KPC cells, but not EL4 cells, led to a significant reduction in myotube diameter. Interestingly, myotube diameter did not significantly reduce in Fn14 knockdown cultures compared to control siRNA cultures in response to treatment with KPC CM (**Fig. 3E, F**). In a separate experiment, we confirmed that Fn14 siRNA drastically reduced protein levels of Fn14 in cultured myotubes (**Fig. 3G, H**). These results suggest that signaling from Fn14 receptor mediates the loss of skeletal muscle mass in response to KPC tumor-derived factors both *in vivo* and *in vitro*.

### Fn14 activates the UPR in skeletal muscle of tumor-bearing mice

Previous studies have demonstrated that TWEAK-Fn14 system induces the activation of ubiquitin-proteasome system including the gene expression of MAFBx and MuRF1, two established markers of the ubiquitin proteasome system. Analysis of skeletal muscle of control and KPC tumor-bearing Fn14^fl/fl^ and Fn14^mKO^ mice revealed that there was a considerable increase in the mRNA levels of MAFbx and MuRF1 in TA muscle of KPC tumor-bearing Fn14^fl/fl^ mice, but not in Fn14^mKO^ mice compared to corresponding control mice injected with PBS alone (**Fig. 4A, B**).

**FIGURE 4.**
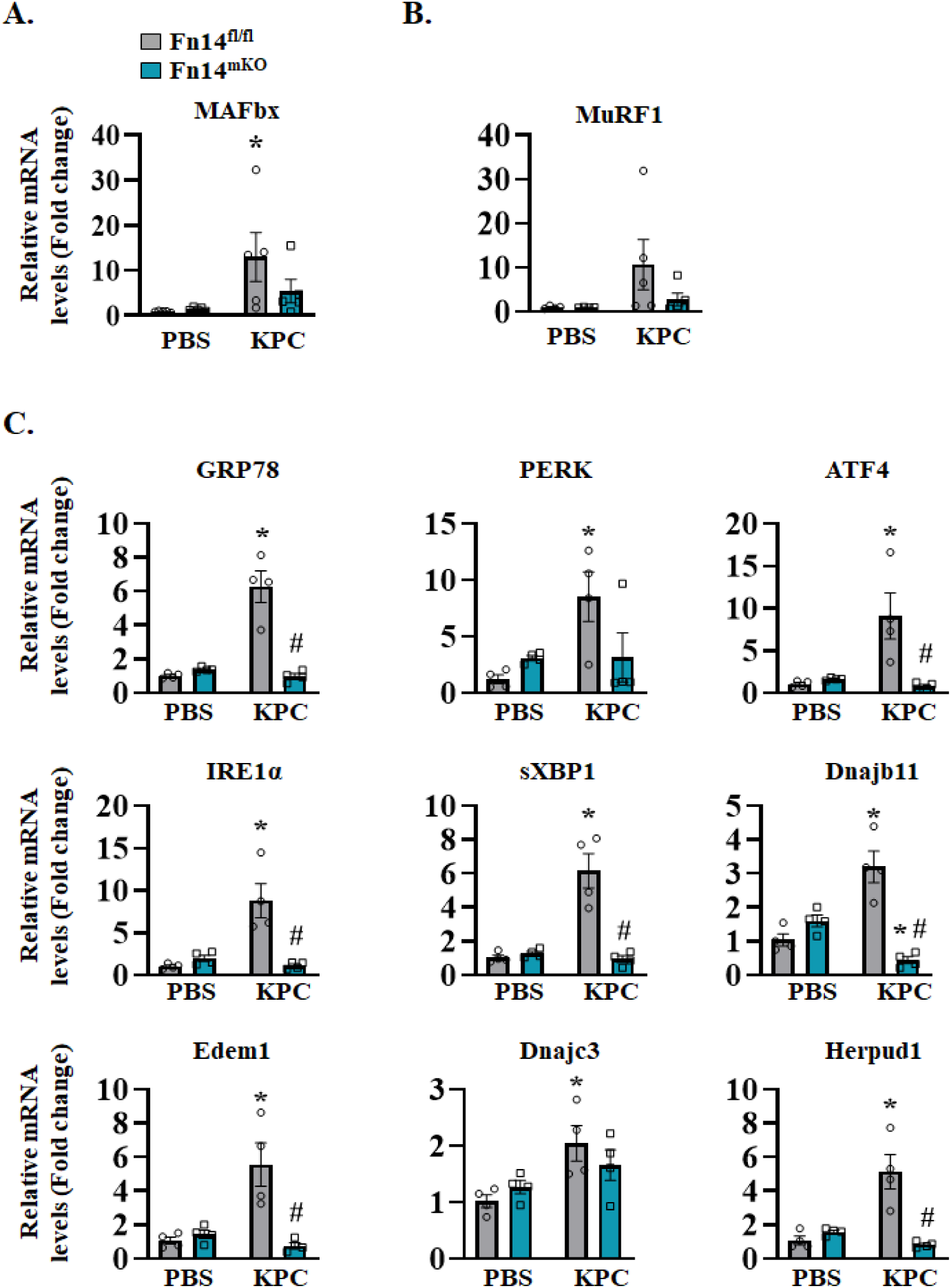
Targeted ablation of Fn14 inhibits the markers of ER stress/UPR in skeletal muscle of KPC tumor-bearing mice. Relative mRNA levels of **(A)** MAFbx, **(B)** MuRF1, and **(C)** GRP78, PERK, ATG4, IRE1α, sXBP1, Dnajb11, Edem1, Dnajc3, and Herpud1 in GA muscle of control and KPC tumor-bearing Fn14^fl/fl^ and Fn14^mKO^ mice. n = 4-5 mice in each group. All data are presented as mean ± SEM and analyzed by two-way ANOVA followed by Tukey’s multiple comparison test. *p ≤ .05, values significantly different from corresponding muscle of control mice injected with PBS alone. ^#^p ≤ 0.05, values significantly different from KPC tumor-bearing Fn14^fl/fl^ mice.

We have previously reported that the markers of ER stress/UPR are highly elevated in skeletal muscle of mouse models of cancer cachexia (15, 37). We investigated whether TWEAK-Fn14 dyad has any role in the regulation of the UPR in skeletal muscle of KPC tumor-bearing mice. Interestingly, we found that mRNA levels of multiple markers of the UPR (i.e., GRP78, ATF4, IRE1α, sXBP1, Dnajb11, Edem1, Dnajc3, and Herpud1) were signficantly reduced in TA muscle of KPC tumor-bearing Fn14^mKO^ mice compared with corresponding KPC tumor-bearing Fn14^fl/fl^ mice (**Fig. 4C**). These results provide intial evidence that the TWEAK-Fn14 system activates the UPR in skeletal muscle during cancer cachexia.

### The TWEAK-Fn14 system induces myotube atrophy through the activation of UPR

While we found reduced expression of multiple components of the UPR in skeletal muscle of KPC tumor-bearing Fn14^mKO^ mice compared to Fn14^fl/fl^ mice, it was not clear whether the TWEAK/Fn14 signaling directly regulates the UPR. To address this issue, cultured mouse primary myotubes were treated with recombinant TWEAK protein for 24 h followed by performing Western blot for a few markers of the UPR. Interestingly, TWEAK increased the levels of phosphorylated PERK and eIF2α (downstream phosphorylation target of PERK) in cultured myotubes (**Fig. 5A**). In addition, protein levels of ATF4, a downstream target of the PERK-eIF2α arm of the UPR (12, 38, 39), were also increased in TWEAK-treated cultures (**Fig. 5A**). Activated IRE1α through its endonuclease activity causes alternative splicing of XBP1 mRNA resulting in the formation of transcriptionally active spliced XBP1 (sXBP1) protein (12, 38, 39). We found that TWEAK also increased the levels of sXBP1 in cultured myotubes (**Fig. 5A**). One of the consequences of the activation of ER stress-induced UPR is the repression of general protein synthesis. While TWEAK has been shown to induce protein degradation, it remains unknown whether TWEAK affects protein synthesis in skeletal muscle. By performing SUnSET assay, we measured the effect of TWEAK on the rate of protein synthesis in cultured mouse primary myotubes. Interestingly, recombinant TWEAK protein repressed the rate of protein synthesis in cultured myotubes in a dose-dependent manner (**Fig. 5B, C**). Moreover, our results showed that TWEAK represses protein synthesis through its bona fide receptor Fn14 because siRNA-mediated knockdown of Fn14 improved protein synthesis and prevented the increase in sXBP1 protein in TWEAK-treated cultured myotubes (**Fig. 5D, E**).

**FIGURE 5.**
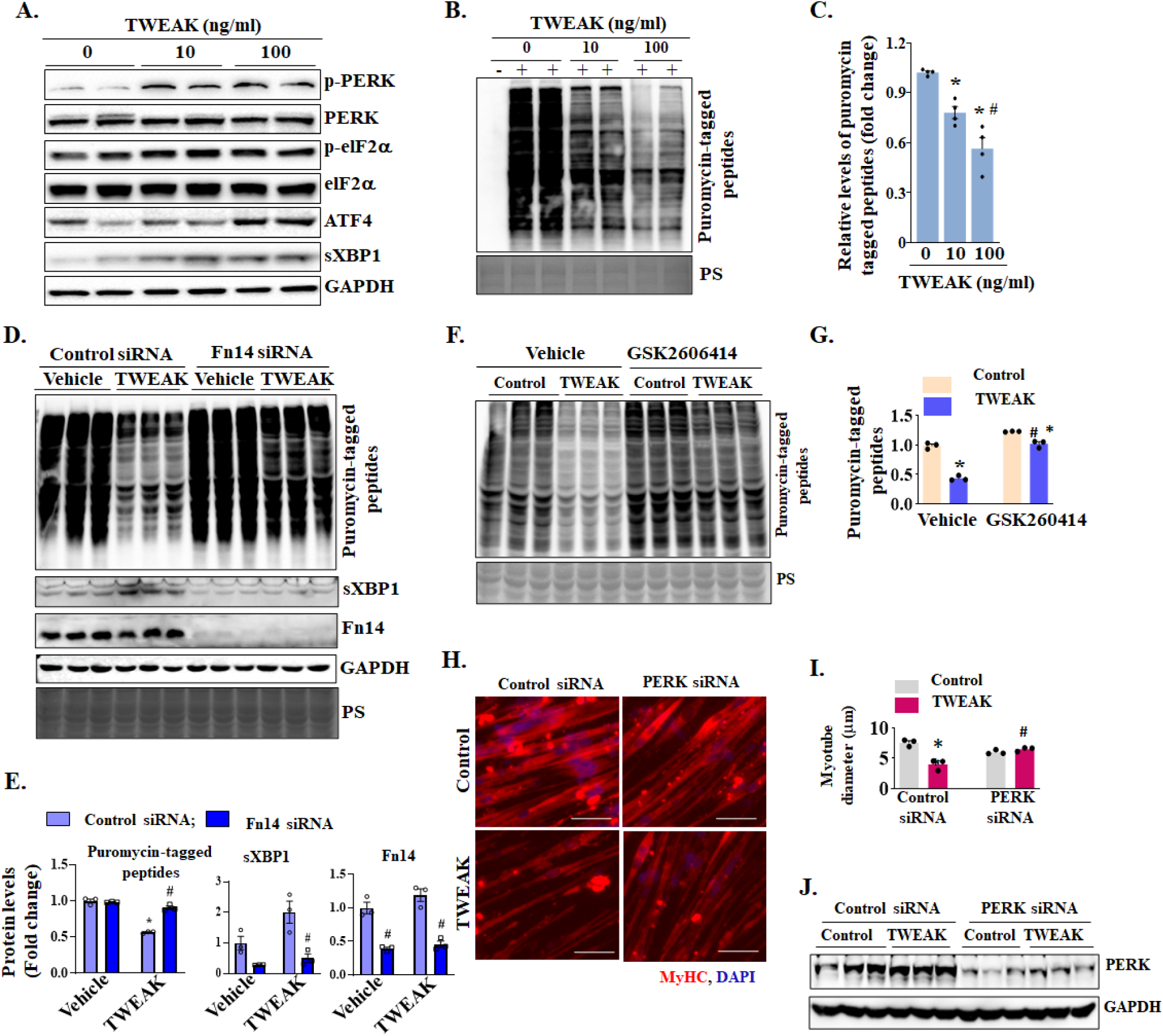
TWEAK inhibits protein synthesis and induces myotube atrophy through the activation of the UPR. **(A)** Representative immunoblots of p-PERK and total PERK, p-eIF2α and total eIF2α, total ATF4 and spliced XBP1 (sXBP1) protein in control and TWEAK-treated cultured primary myotubes. Primary myotubes were treated with indicated concentration of recombinant TWEAK protein for 24h followed by incubation with 1 µM puromycin for 30 minutes before collection **(B)** Representative immunoblot, and **(C)** densitometry analysis showing puromycin-tagged peptides. n = 4 biological replicates in each group. *p ≤ 0.05, vs. 0 ng/ml TWEAK cultures. #p ≤ 0.05, vs. 10ng/ml TWEAK-treated cultures analyzed by one-way ANOVA. Primary myotubes were transfected with control or Fn14 siRNA for 24 h, followed by treatment with vehicle alone (i.e., PBS) or 100 ng/ml TWEAK for an additional 24 h. The cultures were also treated with 1 µM puromycin for 30 minutes before collection**. (D)** Representative immunoblot, and **(E)** densitometry analysis of the levels of puromycin-tagged peptides, sXBP1, and Fn14 protein in control and TWEAK-treated cultures. n = 3 in each group. *p ≤ 0.05, vs. control siRNA transfected cultures treated with vehicle alone. #p ≤ 0.05, vs. corresponding control siRNA transfected cultures. Myotubes were treated with vehicle alone (i.e., DMSO) or 1 μM GSK2606414 for 24 h followed by treatment with 100 ng/ml TWEAK for an additional 24 hours. Cells were incubated with 1 µM puromycin for 30 minutes before collection. **(F**) Representative immunoblot and **(G)** densitometry analysis of puromycin-tagged peptides in control and 100 ng/ml TWEAK-treated primary myotubes incubated without or with PERK inhibitor, GSK2606414. *p ≤ 0.05, vs. control cultures incubated without TWEAK and without GSK2606414. #p ≤ 0.05, significantly different from TWEAK-treated cultures incubated without GSK2606414. **(H)** Representative images of control or PERK siRNA transfected myotubes treated with 100 ng/ml TWEAK or vehicle immunostained for MyHC. Nuclei were counterstained with DAPI. Scale bars, 50 μm. **(I)** Quantification of average myotube diameters of control or PERK siRNA transfected myotubes treated with 100 ng/ml TWEAK or vehicle alone. n = 3 biological replicates in each group. *p ≤ 0.05, values significantly different from control culture transfected with control siRNA; ^#^p ≤ 0.05, values significantly different from TWEAK-treated cultures transfected with control siRNA. **(J)** Representative immunoblot of PERK and unrelated protein GAPDH in control and 100 ng/ml TWEAK-treated primary myotubes transfected with control siRNA or *PERK* siRNA. n = 3 biological replicates in each group. All data are presented as mean ± SEM and analyzed by one-way or two-way ANOVA followed by Tukey’s multiple comparison test.

In another experiment, we investigated whether TWEAK represses protein synthesis through the activation of the PERK arm of the UPR. Results showed that pretreatment of myotubes with GSK2606414, a highly specific inhibitor of PERK (40, 41), restored protein synthesis in TWEAK-treated myotubes (**Fig. 5F, G**). We next sought to determine whether PERK has a role in TWEAK-induced myotube atrophy. Primary myotubes were transfected with control or PERK siRNA for 24 h. The myotubes were then treated with recombinant TWEAK protein and after 48 h the cultures were fixed and immunostatined for MyHC (**Fig. 5H**). Interestingly, we found that siRNA-mediated knockdown of PERK signficantly increased average myotube diameter in TWEAK-treated cultures (**Fig. 5H-J**). Collectively, these results suggest that TWEAK/Fn14 inhibits protein systhesis and causes myotube atrophy through the activation of the UPR.

### Fn14 knockdown attenuates tumor growth *in vivo*

TWEAK and Fn14 are overexpressed and linked to growth and metastases of various tumors, such as gliomas, prostate, and breast cancers (19–22). There is also a published report suggesting that Fn14 signaling in tumor cells can contribute to muscle wasting in C26 mouse model of cancer cachexia (33). We investigated the role of Fn14 signaling in KPC cells on tumor growth in mice. For this experiment, cultured KPC cells were transduced with lentiviral particles expressing a scrambled shRNA or Fn14 shRNA along with lentiviral particles expressing firefly luciferase. Mice injected with PBS served as control. After 48 h, knockdown of Fn14 in KPC cells was confirmed by performing western blot (**Fig. 6A, B**). Next, control or Fn14 shRNA expressing KPC cells were injected in pancreas of 10-week old wild-type mice. Tumor growth was monitored by bioluminescence imaging. There was a noticeable reduction in signal intensities in mice transplanted with Fn14 shRNA expressing KPC cells compared with those transplanted with scrambled shRNA-expressing KPC cells at day 26 (**Fig. 6C**). The four-paw grip strength of KPC tumor-bearing mice was significantly reduced compared to control mice. However, there was no significant difference in the grip strength of the mice injected with scrambled or Fn14 knockdown KPC cells (**Fig. 6D**). Next, the mice were euthanized, and tumor and hindlimb muscles were isolated and weighed. Consistent with bioluminescence imaging results, wet weight of Fn14 shRNA-expressing KPC tumor was significantly lower compared to scrambled shRNA-expressing tumors (**Fig. 6E, F**). Western blot analysis confirmed that the levels of Fn14 protein were significantly reduced in Fn14 knockdown KPC tumors compared to scrambled shRNA-expressing KPC tumors (**Fig. 6G, H**).

**FIGURE 6.**
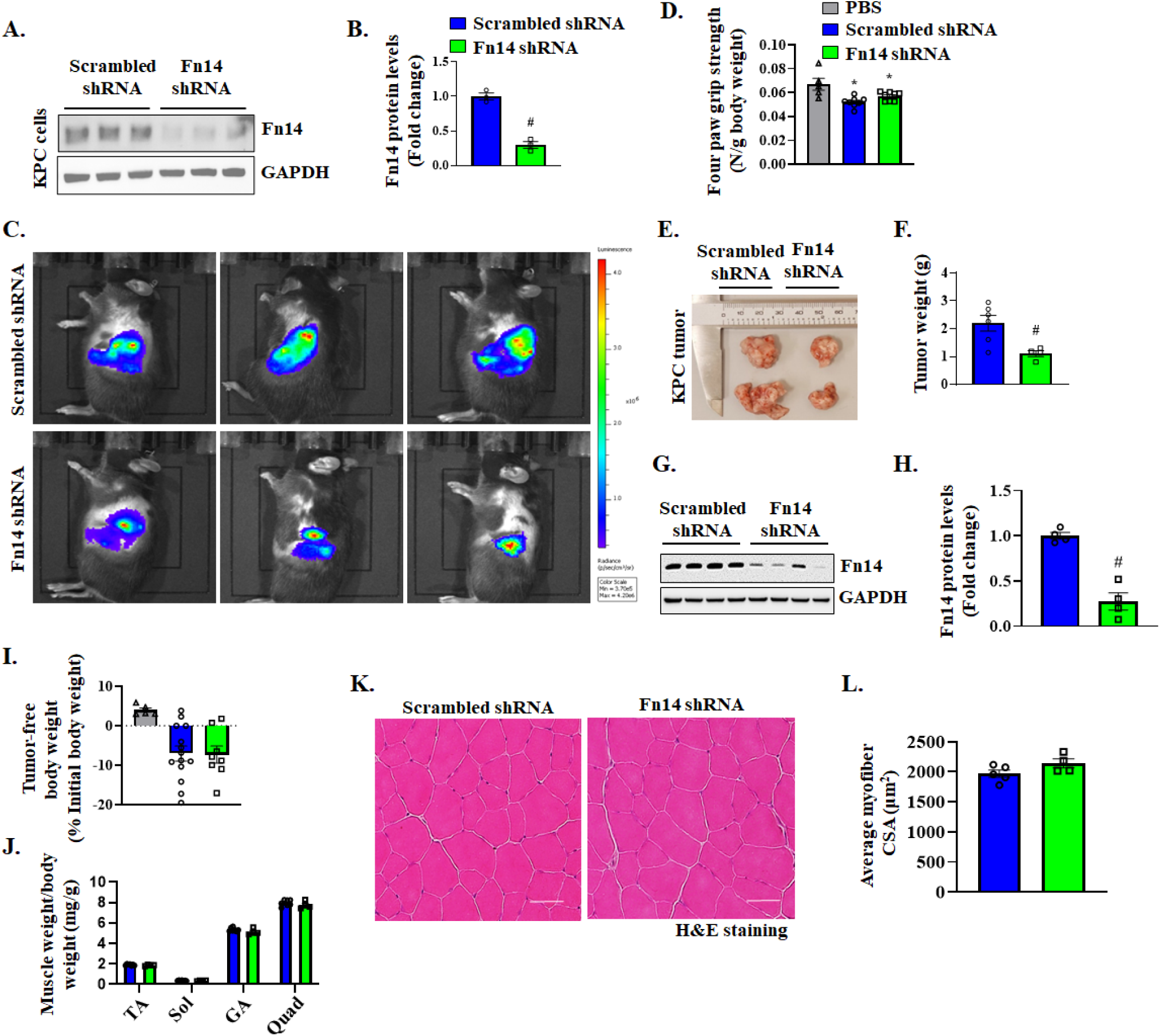
Knockdown of Fn14 inhibits KPC tumor growth in mice. KPC cells were transduced with a scrambled shRNA or Fn14 shRNA and Renilla luciferase-expressing lentiviral particles for 36 h. **(A)** Representative immunoblots, and **(B)** densitometry analysis of Fn14 protein levels in scrambled and Fn14 shRNA KPC cells. 10-week old male C57BL/6J mice were injected with 2 x 10^5^ scrambled shRNA- or Fn14 shRNA-expressing KPC cells in the pancreas and monitored for 21 days. **(C)** Representative bioluminescence images of KPC tumors in mice inoculated with control or Fn14 knockdown KPC cells on day 21. **(D)** Quantification of average four-paw grip strength normalized by body weight of mice injected with PBS alone or scrambled shRNA- or Fn14 shRNA-expressing KPC cells. n = 5-8 mice in each group. **(E)** Representative picture of tumors excised from mice injected with control or Fn14 shRNA-expressing KPC cells. **(F)** Wet weight of control and Fn14 knockdown KPC tumor in mice. n = 4-6 mice in each group. **(G)** Representative immunoblot, and **(H)** densitometry analysis of Fn14 protein in Scrambled shRNA and Fn14 shRNA-expressing KPC tumors. **(I)** Percentage change in tumor free body weight of mice injected with PBS alone or scrambled shRNA or Fn14 shRNA expressing KPC cells. **(J)** Average wet weight of Tibialis anterior (TA), Soleus (Sol), gastrocnemius (GA), and quadriceps (Quad) muscle of control and Fn14 knocked down KPC tumor-bearing mice. **(K)** H&E-stained sections of the TA muscle of control and Fn14 knockdown KPC tumor-bearing mice. Scale bars, 50 μm. **(L)** Quantification of average myofiber CSA in TA muscle of control and Fn14 knockdown KPC tumor-bearing mice. n = 4-5 mice in each group. All data are presented as mean ± SEM and analyzed by one-way ANOVA followed by Tukey’s multiple comparison test or unpaired Student *t* test. *p ≤ 0.05, values significantly different from PBS corresponding group. ^#^p ≤ 0.05, values significantly different from scrambled shRNA KPC-injected mice.

Our analysis also showed that while body weight was reduced in KPC tumor-bearing mice compared to mice injected with PBS, there was no significant difference in tumor-free body weight between scrambled and Fn14 knockdown KPC tumor-bearing mice (**Fig. 6I**). Similarly, there was no significant difference in the wet weight of TA, soleus, GA, or quadriceps (QUAD) muscle of the scrambled or Fn14 knockdown KPC tumor-bearing mice (**Fig. 6J**). We next generated TA muscle transverse sections followed by performing H&E staining and morphometric analysis. Results showed that there was a trend towards increase in average myofiber CSA in TA muscle of mice implanted with Fn14 shRNA-expressing KPC cells compared to scrambled shRNA expressing cells. However, this effect was not statistically significant (**Fig. 6K, L**). Collectively, these results suggest that while silencing Fn14 attenuates KPC tumor growth *in vivo*, it does not have any significant effect on the loss of muscle mass in response to KPC tumor growth.

### The silencing of Fn14 inhibits the proliferation, migration, and invasiveness of KPC cells

To understand the mechanisms by which Fn14 signaling in KPC cells promotes tumor growth *in vivo*, we first investigated the effect of knockdown of Fn14 on KPC cell growth. KPC cells expressing scrambled (control) or Fn14 shRNA were seeded at equal number and the total number of cells were counted at 24 h and 48 h later. The number of cells was significantly reduced in Fn14 knockdown cultures compared to controls at both the time points, suggesting that Fn14-mediated signaling significantly promotes the growth of KPC cells (**Fig. 7A**). To further access the role of Fn14 signaling in the proliferation of KPC cells, we also performed an EdU incorporation assay. For this experiment, KPC cells expressing scrambled or Fn14 shRNA were seeded at equal density, followed by pulse labeling with EdU for 15 min and detection of EdU-positive nuclei (**Fig. 7B**). Results showed that the proportion of EdU-positive cells was significantly reduced in Fn14 knockdown KPC cultures compared to corresponding control cultures (**Fig. 7C**). By performing Annexin V staining followed by FACS analysis, we next investigated whether knockdown of Fn14 affects the apoptosis in KPC cells. However, no difference was observed in the percentage of Annexin V positive KPC cells between cultures expressing either scrambled or Fn14 shRNA (**Figure S1**) suggesting that silencing of Fn14 does not affect the viability of KPC cells.

**FIGURE 7.**
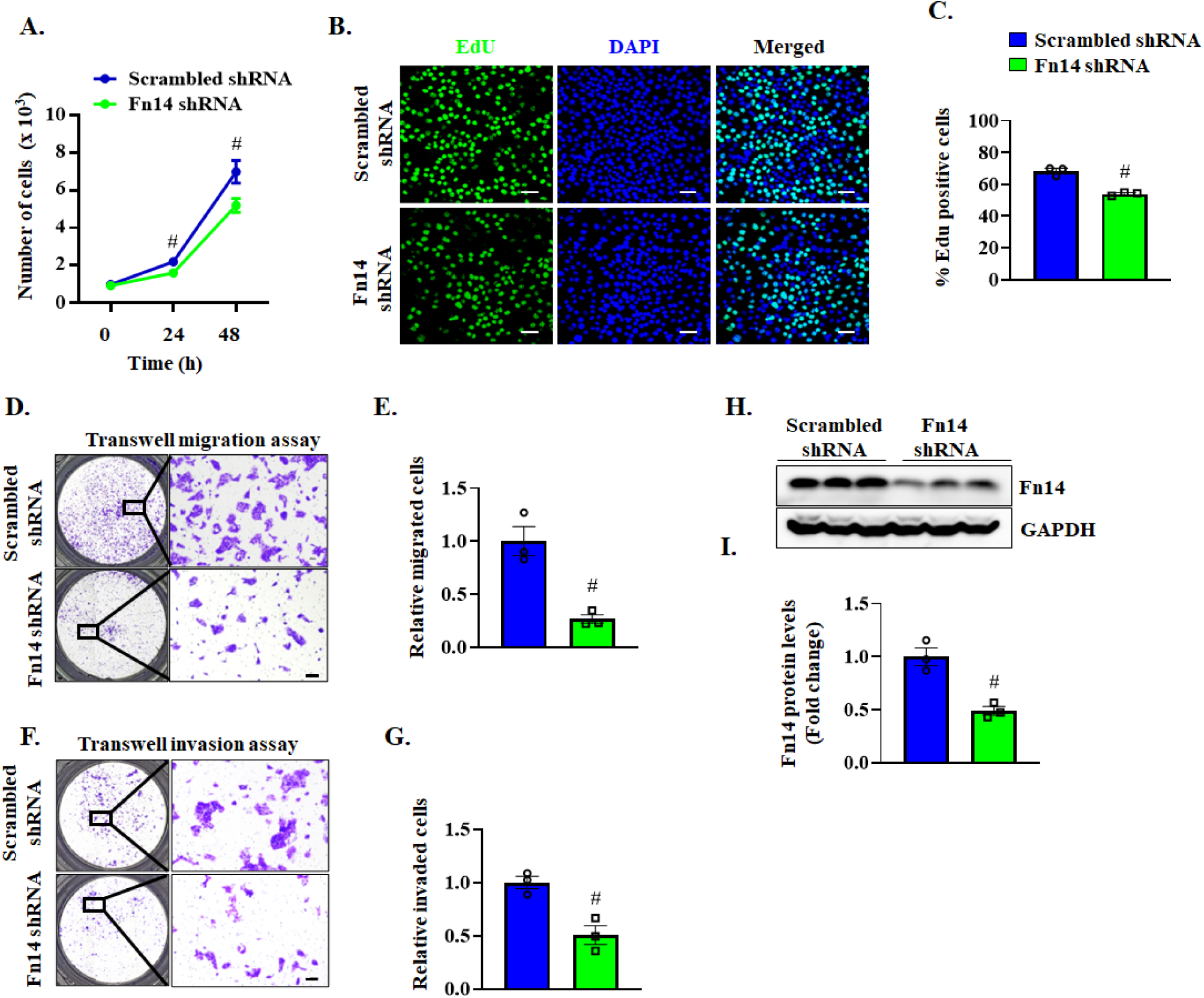
Fn14 promotes proliferation, migration and invasion of KPC cells. **(A)** Cell proliferation was assessed by seeding 1,000 cells per well in a 96-well plate. The graph illustrates the average cell count at each time point, demonstrating cell growth over time. Data are presented as the mean ± 95% confidence interval. **(B)** Representative EdU and DAPI stained images of scrambled or Fn14 shRNA expressing KPC cultures. Scale bar, 50 µm. **(C)** Quantification of EdU-positive cells in scrambled or Fn14 shRNA expressing KPC cultures. Data are presented as the mean ± SEM. n = 3 biological replicates in each group. **(D)** Representative images of migrated cells in transwell migration assay. Scale bars, 50 µm. **(E)** Quantification of the relative migration of control and Fn14 knockdown KPC cells. **(F)** Representative images of invaded cells in transwell invasion assay. Scale bars, 50 µm. **(G)** Quantification of the relative invasive capacity of control and Fn14 knockdown KPC cells. **(H)** Western blot analysis and **(I)** quantification of Fn14 protein in KPC cells transduced with scrambled shRNA or Fn14 shRNA. All data are presented as mean ± SEM and analyzed by unpaired Student *t* test. #p ≤ 0.05, values significantly different from scrambled shRNA expressing cultures.

We next investigated the role of Fn14 in cell migration using a wound healing assay. Control and Fn14 knockdown KPC cells were plated at full confluence, a scratch in a cell monolayer was made, and the images of scratch were captured at a regular interval during cell migration. The scratch gap was significantly higher in Fn14 knockdown cultures compared to controls suggesting that silencing of Fn14 inhibits migration of KPC cells *in vitro* (**Figure S2**). We also employed a transwell assay to study the role of Fn14 in the migration of KPC cells. Consistent with wound healing assay, knockdown of Fn14 significantly reduced the migration of KPC cells in transwell assay (**Fig 7D, E**). Using matrigel coated transwell plates we next measured the effects of knockdown of Fn14 on the invasive capacity of KPC cells. Results showed that knockdown of Fn14 significantly reduced the KPC cell invasion (**Fig. 7F, G**). Western blot analysis confirmed reduced levels of Fn14 protein in Fn14 shRNA expressing KPC cells (**Fig. 7H, I**). Collectively, these results suggest that Fn14 promotes proliferation, migration, and invasiveness of KPC cells.

### Targeted ablation of Fn14 inhibits muscle wasting in the LLC model of cancer cachexia

We also investigated whether muscle-specific ablation of Fn14 can attenuate the loss of muscle mass in other models of cancer cachexia, such as Lewis lung carcinoma (LLC). For this experiment, 10-week old Fn14^fl/fl^ and Fn14^mKO^ mice were injected with 2x10^6^ cells in the flanks. Control mice received injections of saline only. Grip strength measurements were performed after 21 days of injection of LLC cells. While there was significant reduction in grip strength (normalized by body weight) of LLC tumor-bearing Fn14^fl/fl^ mice, there was no reduction in the grip strength of Fn14^mKO^ in response to LLC tumor growth compared to corresponding control mice (**Fig. 8A**). Next, the mice were euthanized, and individual hind limb muscles were isolated and weighed. Intriguingly, there was no significant difference in the wet weight of TA, GA, or QUAD muscle between LLC tumor-bearing Fn14^fl/fl^ and Fn14^mKO^ mice (**Fig. 8B**). We next generated TA muscle transverse sections followed by performing H&E staining, or anti-laminin and DAPI staining (**Fig. 8C, D**). The proportion of myofibers with higher CSA was considerably increased in TA muscle of LLC tumor-bearing Fn14^mKO^ mice compared to Fn14^fl/fl^ mice (**Fig. 8E**). Moreover, average myofiber CSA in TA muscle was significantly higher in LLC tumor-bearing Fn14^mKO^ mice compared to corresponding Fn14^fl/fl^ mice (**Fig. 8F**). Collectively, these results suggest that muscle-specific ablation of Fn14 inhibits muscle atrophy in LLC model of cancer cachexia.

**FIGURE 8.**
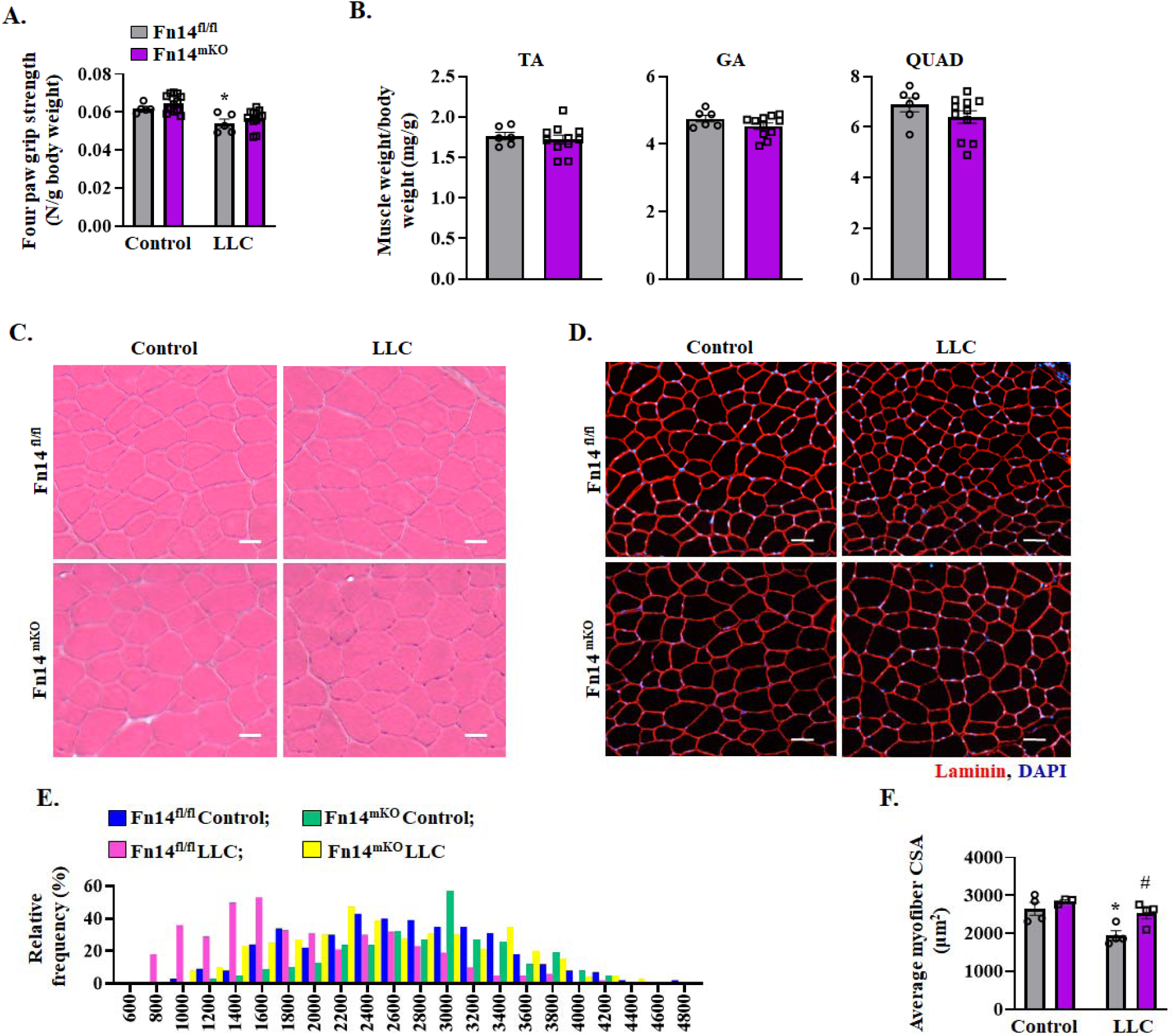
Targeted deletion of Fn14 inhibits LLC tumor-induced muscle wasting. 10-week old Fn14^fl/fl^ and Fn14^mKO^ mice were inoculated with Lewis lung carcinoma (LLC) cells in the dorsal subcutis and monitored for 21 days. **(A)** Quantification of average four-paw grip strength of control and KPC tumor-bearing Fn14^fl/fl^ and Fn14^mKO^ mice normalized by body weight. **(B)** Wet weight of tibialis anterior (TA), gastrocnemius (GA), and quadriceps (Quad) muscle normalized with tumor-free body weight of mice. **(C)** H&E-stained and **(D)** anti-laminin-stained sections of the TA muscle of control and LLC tumor-bearing Fn14^fl/fl^ and Fn14^mKO^ mice. Scale bars, 50 μm. **(E)** Relative frequency distribution of myofiber cross-section area (CSA) in TA muscle of control and LLC tumor-bearing Fn14^fl/fl^ and Fn14^mKO^ mice. **(F)** Average myofiber CSA in control and LLC tumor-bearing Fn14^fl/fl^ and Fn14^mKO^ mice. n = 3-4 mice in each group. All data are presented as mean ± SEM and analyzed by two-way ANOVA followed by Tukey’s multiple comparison test or unpaired Student *t* test. *p ≤ 0.05, values significantly different from corresponding control mice. ^#^p ≤ 0.05, values significantly different from LLC-injected Fn14^fl/fl^ mice.

## DISCUSSION

Loss of skeletal muscle mass is a devastating complication that contributes to both morbidity and mortality of cancer patients (1, 2). Several factors have been identified that may trigger muscle wasting during cancer cachexia (3, 4, 15, 34, 42). Inflammation plays a critical role in skeletal muscle wasting in many chronic disease states, including cancer. Indeed, proinflammatory cytokines, such as TNFα and IL-6 have been suggested to mediate muscle wasting in animal models (3, 4, 43). TWEAK is another proinflammatory cytokine that has been shown to induce muscle wasting in various disuse conditions (25, 29, 30, 32, 44). However, the role of TWEAK - Fn14 system in the regulation of skeletal muscle mass during cancer-induced cachexia remains poorly understood. Our present study demonstrates that the expression of Fn14 is increased in the skeletal muscle of multiple models of cancer cachexia. Furthermore, our results demonstrate that muscle-specific deletion of Fn14 attenuates skeletal muscle wasting in KPC or LLC tumor-bearing mice.

Muscle wasting in many conditions is attributed to enhanced protein degradation due to the increased activation of proteolytic systems, such as the UPS and autophagy (8). Indeed, many catabolic stimuli and tumor-derived factors have been shown to induce the activation of proteolytic systems both *in vivo* and *in vitro* and targeted inhibition of the components of the UPS or autophagy attenuates muscle wasting in animal models (2, 42, 45–49). Furthermore, inhibition in the rate of protein synthesis can also contribute to the loss of skeletal muscle mass during cancer cachexia (50–52). However, the mechanisms leading to the repression of protein synthesis in skeletal muscle of cancer patients remain poorly understood.

ER stress-induced UPR pathways play an important role in the regulation of muscle mass in different conditions (10, 11, 53). It is now increasingly clear that physiological levels of activation of the UPR are important for maintenance of muscle health (14, 54). However, chronic activation of ER stress and UPR can cause deleterious consequences in skeletal muscle (10, 11, 55). Our previous studies have shown that multiple markers of ER stress/UPR are highly elevated in skeletal muscle of mouse models of cancer cachexia (14, 15). Although the mechanisms leading to the activation of the UPR during cancer-induced muscle wasting remain unknown, our studies demonstrated that forced activation of IRE1α or sXBP1 causes myotube atrophy and targeted deletion of XBP1 attenuates muscle wasting in the LLC-tumor-bearing mice (15). Interestingly, in the present study we found growth of KPC tumor in mice also increases the expression of various components of the PERK and IRE1α/XBP1 arms of the UPR. More importantly, markers of the UPR as well as the UPS are significantly reduced in skeletal muscle of KPC tumor-bearing muscle-specific Fn14 knockout mice (**Fig. 4B**) suggesting that in addition to proteolytic systems, the TWEAK-Fn14 system may be inducing muscle wasting during cancer cachexia through the activation of the UPR. Furthermore, it is also plausible that the components of the UPR crosstalk with proteolytic systems and other catabolic signaling pathways to induce muscle wasting in tumor-bearing hosts (56, 57).

Repression of general protein synthesis is one of the important consequences of the activation of the PERK arm of the UPR (58). While the TWEAK/Fn14 signaling has been shown to activate proteolytic pathways in cultured myotubes and animal models (26, 44, 59), the role of this ligand-receptor dyad in protein synthesis in skeletal muscle is unknown. Our experiments in the present study demonstrate that TWEAK strongly inhibits protein synthesis in cultured primary myotubes. Moreover, recombinant TWEAK protein activates the components of PERK and IRE1α/XBP1 signaling in cultured myotubes. Our experiments also suggest that the activation of PERK is the main mechanism by which TWEAK represses protein synthesis and causes atrophy in cultured myotubes (**Fig. 5**). These results are consistent with our findings *in vivo* demonstrating that muscle-specific deletion of Fn14 inhibits the gene expression of markers of both the PERK and IRE1/XBP1 arms of the UPR (**Fig. 4**). However, the signaling mechanisms by which TWEAK/Fn14 system activates the UPR in skeletal muscle remain unknown and should be a focus of future investigations.

Previous studies have shown that TWEAK/Fn14 system also plays an important role in growth and metastasis of multiple tumor types (23, 60). However, the role of Fn14 in the regulation of cachectic pancreatic cancer growth remained unknown. Our results demonstrate that silencing of Fn14 significantly reduces the growth of KPC tumor in the pancreas of mice (**Fig. 6**). Furthermore, our experiments suggest that inhibition of Fn14 signaling reduces the proliferation, migration and invasiveness of KPC cells (**Fig. 7**). These results are consistent with previously published reports also suggesting the importance of TWEAK/Fn14 signaling in cell proliferation, migration, and angiogenesis (19–22). Indeed, TWEAK/Fn14 axis has been shown to activate NF-κB and MAPK signaling which promotes cell proliferation and migration (18, 61). While we found that silencing of the Fn14 attenuated KPC tumor growth, surprisingly, it did not have any significant effect on muscle wasting in mice (**Fig. 6J-L**). This could be attributed to the fact that knockdown of Fn14 did not completely prevent the tumor growth and the size of the Fn14-knocked down KPC tumor was sufficient to release cachectic factors and activate host-derived mechanisms to induce muscle wasting. Nevertheless, the inhibition of KPC tumor growth upon silencing of Fn14 suggests that blocking TWEAK/Fn14 signaling can be an important approach to attenuate muscle wasting and growth of tumor in pancreatic cancer patients.

Johnston et al., (33) previously reported that Fn14 expression in tumors, but not in skeletal muscle, contributes to muscle wasting during cancer-induced cachexia. In this study, the authors showed that overexpression of Fn14 in tumor cells increases tumor growth and severity of cachexia, the latter of which was rescued by injecting the mice with their newly developed Fn14 neutralizing antibodies at the onset of cachexia. Additionally, the study showed that anti-Fn14 treatment also inhibits cachexia in C26 tumor model, despite having no significant effect on tumor size over the duration of the study. By contrast, our study showed that knockdown of Fn14 attenuates the growth of KPC tumor in mice, yet it leads to a similar degree of muscle wasting (**Fig. 6J-L**). Since different tumor types can induce cachexia through distinct mechanisms, it is possible that in some other types of cancers, including C26 model, the TWEAK/Fn14 signaling in tumors will also contribute to muscle wasting. Another possibility of this discrepancy might stem from the approaches of targeting Fn14 activity, i.e., kinetic distribution of Fn14 antibodies in mice versus genetically modified cells used in our study. Since the neutralizing Fn14 antibodies were injected intraperitoneal, the anti-Fn14 might target both tumor and host tissues, including muscle, which can significantly alter tumor growth and progression of cachexia. Furthermore, the authors observed no significant differences in cancer-induced muscle wasting in TWEAK and Fn14-global knockout mice, suggesting that TWEAK/Fn14 system in host tissues does not promote cachexia (33). However, global deletion of TWEAK or Fn14 might change the entire cytokine milieu, immune profile and metabolic states of different tissues (18, 30), that can have a confounding effect on cachexia. Interestingly, a recent study by Liu et al., (62) identified a novel regulatory circuitry in which macrophages facilitate TWEAK secretion from pancreatic tumor cells, that eventually induce muscle wasting through Fn14 signaling. Consistent with this study (62), our results provide genetic evidence that deletion of Fn14 in myofibers attenuates cancer-induced muscle wasting in KPC and LLC mouse models of cancer cachexia.

In summary, our study demonstrates that TWEAK/Fn14 signaling in myofibers mediates skeletal muscle wasting in animal models of cancer cachexia. Furthermore, our experiments indicate that blocking of Fn14-mediated signaling can inhibit growth of pancreatic cancer *in vivo*. While more investigations are needed, the present study provides initial evidence that the TWEAK/Fn14 signaling is activated in skeletal muscle during cancer cachexia and inhibition of this pathway could be a potential approach to prevent muscle wasting in cancer patients.

## MATERIALS AND METHODS

### Animals

C57BL/6J mice were purchased from The Jackson Laboratory. Floxed Fn14 (i.e., Fn14^fl/fl^) mice as described (63) were provided by Dr. Linda Burkly of Biogen, Indc. Fn14^fl/fl^ mice were crossed with muscle creatine kinase (MCK)-Cre (Strain: B6.FVB(129S4)-Tg(Ckmm-cre)5Khn/J, Jackson Laboratory, Bar Harbor, ME) mice to generate muscle-specific Fn14 knockout (henceforth Fn14^mKO^) mice and littermate Fn14^fl/fl^ mice (32). The KPC cell orthotopic xenograft was conducted following a protocol as described (34). Briefly, 2 × 10^6^ KPC cells (resuspended in 20 µL Phosphate buffered saline [PBS] containing 50% Matrigel or equal volume of PBS containing 50% Matrigel as control) were injected into the tail of the pancreas of 10-wk-old male C57BL/6J mice under anesthesia. The openings in muscle and skin were sutured or sealed by wound clips, respectively. The KPC tumor growth was monitored using *in vivo* bioluminescence imaging using an IVIS Lumina XR *in vivo* imaging system (Caliper, Mountain View, CA). Briefly, mice were given intraperitoneal injection of firefly Luciferin (150 mg/kg; PerkinElmer). After 10 minutes, the mice were anesthetized using 3% isoflurane, put in the IVIS imaging box, and imaged laterally. Tumor size was observed by normalized luminescence acquisition (IVIS software). The LLC cell subcutaneous xenograft in 10-wk-old C57BL/6J mice was established using a protocol as previously described (14, 15). Development of cachexia was monitored by body weight and four paws grip strength. Mice were euthanized and the samples were analyzed after cachexia had been established. All experiments were approved by the Animal Care Operations and the Institutional Animal Care and Use Committee at the University of Houston.

### Grip strength test

Total 4-paw grip strength of mice were measured using a digital grip-strength meter (Columbus Instruments) and normalized by body weight as described (15).

### Histology and morphometric analysis

Individual TA and soleus muscles were isolated from mice, snap-frozen in liquid nitrogen, and sectioned with a microtome cryostat. For the assessment of muscle morphology, 10-μm-thick transverse sections of TA and soleus muscle were stained with hematoxylin and eosin (H&E) dye. Muscle sections were also processed for immunostaining for laminin protein to mark the boundaries of myofibers. Briefly, frozen TA and soleus muscle sections were fixed in acetone, blocked in 2% bovine serum PBS for 1 h, and incubated with rabbit anti-laminin in blocking solution at 4°C overnight under humidified conditions. The sections were washed briefly with PBS before incubation with goat anti-rabbit Alexa Fluor 468-conjugated secondary antibody for 1 h at room temperature and were then washed three times for 15 min each time with PBS. All stained and immunofluorescence sections were visualized, and images captured using an inverted microscope (Nikon Eclipse Ti-2E Inverted Microscope), a digital camera (Digital Sight DS-Fi3, Nikon) and NIS Elements AR software (Nikon). The NIH ImageJ software was used for quantitative analysis and image levels were equally adjusted using Photoshop CS6 software (Adobe).

### Generation of Fn14 shRNA expressing lentiviral particles

The target siRNA sequences for mouse Fn14 mRNA were identified using BLOCK-iT RNAi Designer online software (Life Technologies). The shRNA oligonucleotides were synthesized to contain the sense strand of target sequences for mouse Fn14 shRNA (GCCAAGCTGGAAGCCATTAAT), a short spacer (CTCGAG), and the reverse complement sequences followed by five thymidine as an RNA polymerase III transcriptional stop signal. Oligonucleotides were annealed and cloned into pLKO.1-Puro plasmid with AgeI/EcoRI sites. The insertion of shRNA sequence in the plasmid was confirmed by DNA sequencing.

### Cell culture

Primary myoblasts were prepared from hind limb muscle of C57BL6 mice following a procedure as described (64). These cells were cultured in growth medium (GM; Dulbecco’s modified Eagle’s medium [DMEM] containing 10% FBS). To induce differentiation, the cells were incubated in differentiation medium (DM; DMEM supplemented with 2% horse serum) for 48 h. The EL4 cell line was purchased from ATCC. KPC cells were kindly provided Elizabeth Jaffee (Johns Hopkins University, Baltimore, MD) (65) and cultured in RPMI1640 supplemented with 10% FBS.

### Preparation of tumor cell conditioned medium

EL4 cells were maintained in suspension in DMEM, supplemented with 10% horse serum (HS) and 1% penicillin-streptomycin (P/S). For conditioned media (CM) preparation, when the cell density reached 1 × 10 cells/mL, the culture media was replaced with fresh media. After 48 hours, the CM were collected, centrifuged, and filtered using 0.45-μm-pore-size filters. KPC cells were cultured in DMEM, supplemented with 10% fetal bovine serum (FBS) and 1% P/S. When the cells reached 80-90% confluence, the culture medium was replaced with fresh media. After 48 hours, the CM were collected, centrifuged, and filtered using 0.45-μm-pore-size filters. The CM was diluted 1:4 with fresh DM for treatment of cultured myotubes and subsequent analysis of myotube atrophy.

### Cell Growth Assay

The effect of knockdown of Fn14 on the proliferation of KPC cells was evaluated following a protocol as described (66). Briefly, cells were seeded at 1,000 cells/well in a 96-well tissue culture plates. The plates were scanned using the EnSight Multimode Plate Reader equipped with well-imaging technology (PerkinElmer, MA, United States). Cell count was obtained by digital phase and brightfield imaging.

### EdU incorporation assay

The Edu incorporation assay was performed using the Click-iT™ Plus EdU Cell Proliferation Kit for Imaging, Alexa Fluor™ 488 dye (Catalog number C10637). Briefly, 5x10^4^ cells were seeded per well of a 12-well plate. After 36 hours, the cells were incubated with 10 µM Edu for 15 min and then fixed with 4% PFA, permeabilized with 0.5% Triton, and then stained with Alexa Fluor 488 dye. The nucleus was visualized with DAPI. The images were taken using Nikon microscopy, and EdU-positive cells were quantified using ImageJ software.

### Wound healing assay

Cultured cells were seeded into 6-well plates at a density of 8 × 10^5^ cells per well. When the monolayer was formed, a scratch of the cell monolayer was made using sterile 1 ml micropipette tip. The closure of the gap was imaged at different time intervals under 4 × field, and the wound closure speed was evaluated using the ImageJ software (NIH).

### Transwell migration and invasion assay

For the Transwell migration assay, 5 × 10^4^ cells in 200 µL serum-free DMEM were reseeded into the top of the insert of a Boyden chamber (Corning Inc., Corning, NY, USA), while 800 µL medium with 10% FBS was loaded into the well below. After 24 h incubation, migrating cells that passed through the filter were stained with 0.1% crystal violet solution. For the Transwell invasion assay, all procedures were similar except that transwell membrane was coated with 100 µl of 300 µg/mL Matrigel (BD Bioscience, San Jose, CA, USA). The cells that passed through the filter were stained with 0.1% crystal violet solution, imaged and the crystal violet signal intensity divided by total area was quantified using the ImageJ software.

### Cell viability assays

For analysis of cell viability, we performed Annexin V staining using the Annexin V Conjugates for Apoptosis Detection Kit following a protocol suggested by the manufacturer (ThermoFisher Scientific).

### Quantitative real-time PCR (QRT-PCR)

Total RNA isolated from GA and QUAD muscle tissues of mice was subjected to reverse transcription and real-time quantitative PCR (qPCR) analysis, as previously described (15). Normalization of data was accomplished using the endogenous control β-actin, and the normalized values were subjected to a threshold cycle (2^−ΔΔ*CT*^) formula to calculate the fold change between the control and experimental groups. The sequence of the primers used for QRT-PCR is described in **Table S1**.

### Surface sensing of translation assay

Protein synthesis was measured by non-isotope-labeled surface sensing of translation (SUnSET), as described (67). Myotubes were treated with TWEAK or other agents followed by addition of 0.1 μM puromycin for 30 min. The cells were collected, protein extracts were made, and newly synthesized protein was detected by Western blot with anti-puromycin (1:1000; EMB Millipore Darmstadt, Germany) as the primary antibody.

### Western blotting

Relative levels of various proteins were determined by performing Western blot analysis as described (15). Quantitative estimation of the intensity of each band was performed using ImageJ software (NIH). Antibodies used for Western blot are described in **Table S2**. The uncropped immunoblot images are presented in Supplemental **Figure S3**.

### Statistical analysis

Results are expressed as means ± SEM. An unpaired, 2-tailed Student’s t test was used to compare quantitative data populations with normal distribution and equal variance, one way or two-way ANOVA followed by Tukey’s multiple comparison test. A value of p ≤ 0.05 was considered significant.

## ACKNOWLEDGEMENTS

This work was supported by National Institute of Health grant AR081487 to AK. We are thankful to Dr. Linda Burkly of Biogen Inc. (Cambridge, MA) for providing floxed Fn14 mice.

