## Supplementary material for "The TWEAK/Fn14 signaling promotes skeletal muscle wasting during cancer cachexia": Figure S1-S3, Tables S1 and S2

### Supplemental Data File

**FIGURE S1**

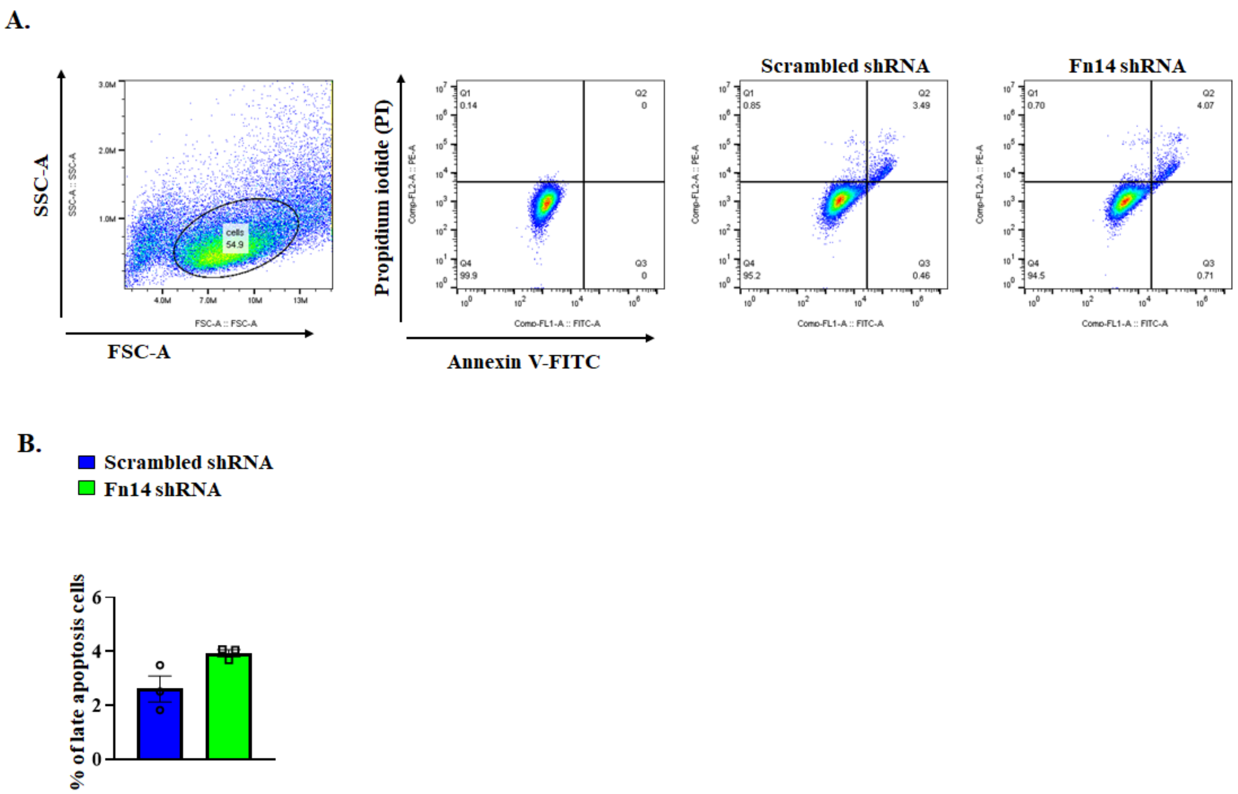


**FIGURE S1. Effect of F14 knockdown on the survival of KPC cells. (A)** Annexin V-FITC staining for apoptotic cells, and **(B)** quantification of late apoptotic cells in KPC cells transduced with scrambled shRNA or Fn14 shRNA. All data are presented as mean ± SEM and analyzed by unpaired Student *t* test. n = 3 biological replicates in each group. No statistically significant differences were obtained by unpaired student t test.

FIGURE S2

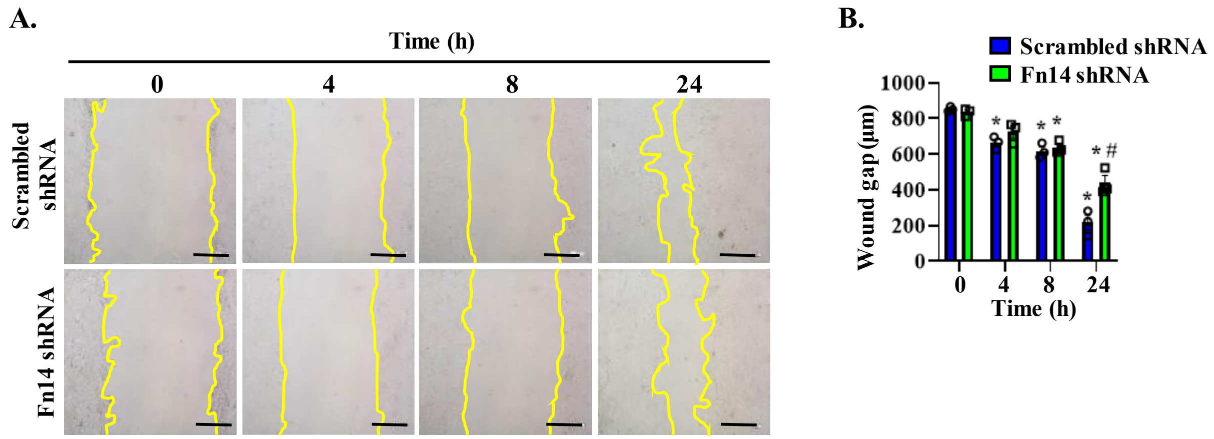


**FIGURE S2. Silencing of Fn14 inhibits KPC cell migration.** Scrambled and Fn14 shRNA-expressing KPC cells were plated at full confluency and a scratch in a cell monolayer was made. **(A)** Representative images, and **(B)** quantification of wound gap at different time points. n = 3 biological replicates in each group. All data are presented as mean ± SEM and analyzed by two-way ANOVA followed by Tukey's multiple comparison test. $^{*}p \leq 0.05$, values significantly different from corresponding cultures at 0 h. $^{\#}p \leq 0.05$, values significantly different from scrambled shRNA-expressing cultures at 24h.

**FIGURE S3**

**Figure 1C**

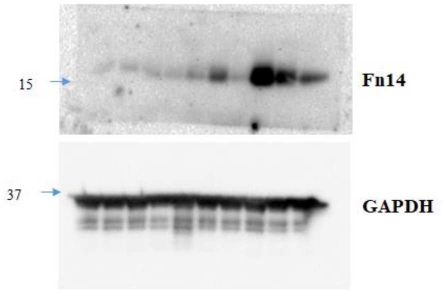


**Figure 3G**

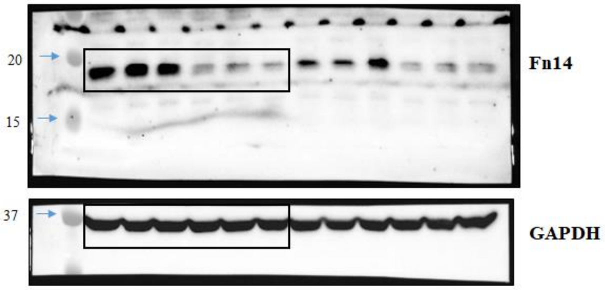


**Figure 5A**

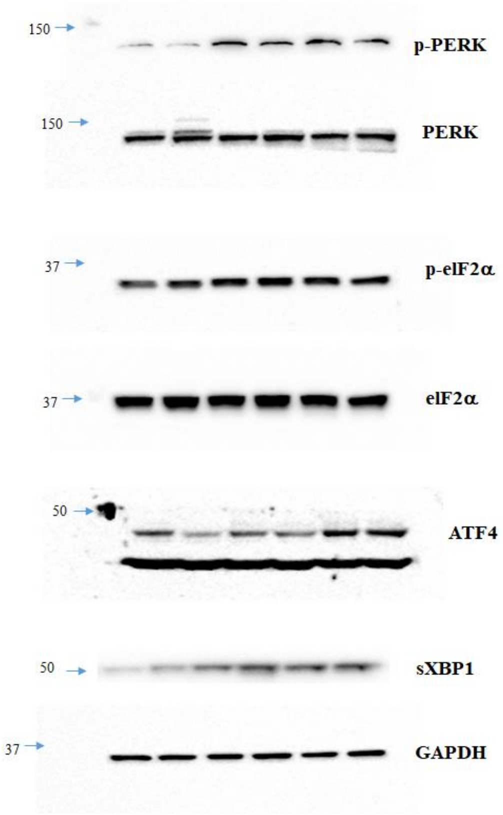

**FIGURE S3 (continuation)**

**Figure 5B**

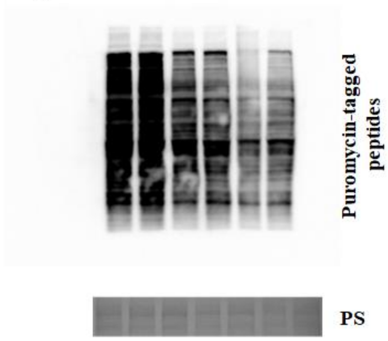


**Figure 5D**

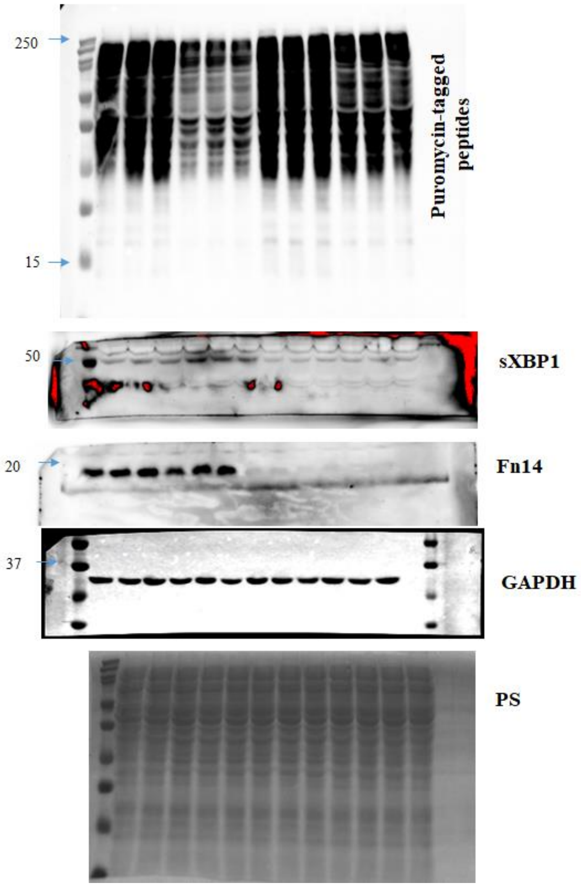

#### FIGURE S3 (continuation)

##### Figure 5F

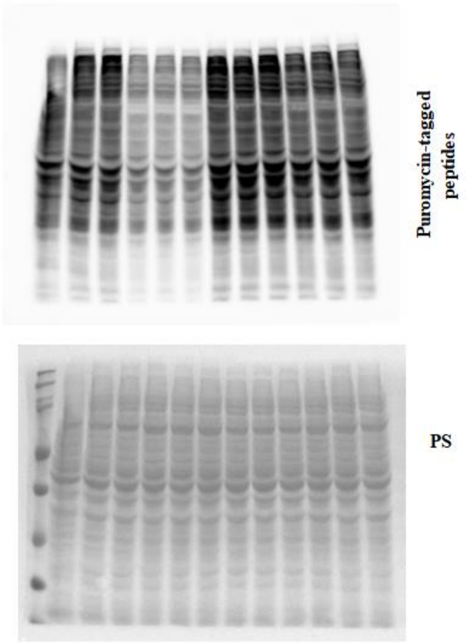


##### Figure 5J

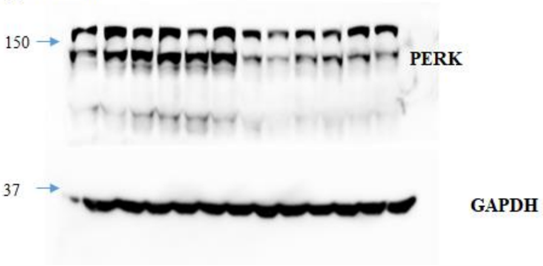


##### Figure 6A

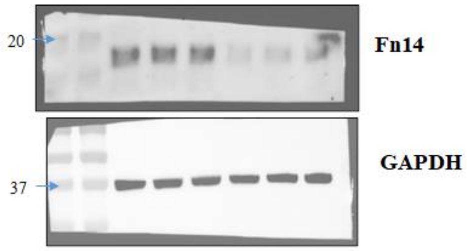


##### Figure 6G

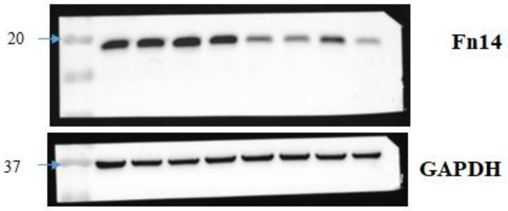


##### Figure 7H

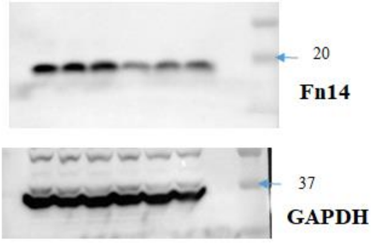


**FIGURE S3. Uncropped western blot gel images used in this study.**

**Table S1. List of primers used for PCR/qRT-PCR analysis.**

| Name | Forward primer (5'-3') | Reverse primer (5'-3') |
|---|---|---|
| TWEAK | GCTACGACCGCCAGATTGGG | GCCAGCACACCGTTCACCAG |
| Fn14 | AAGTGCATGGACTGCGCTTCTT | GGAAACTAGAAACCAGCGCCAA |
| MAFBX | GTCGCAGCCAAGAAGAGAAAGA | TGCTATCAGCTCCAACAGCCTT |
| MURF1 | TACTGCATCTCCATGCTGGTG | TGGCGTAGAGGGTGTCAAACTT |
| GRP78 | CGT GGA GAT CAT AGC CAA CGA T | ATT CCA AGT GCG TCC GAT GA |
| PERK | ACT CCT GTC TTG GTT GGG TCT GAT | CGT GCT CCG CTT ATT CCT TTC T |
| ATF4 | CCTCTTCACGAAATCCAGCAGCA | CCATGAGGTTTCAAGTGCTTG |
| IRE1 | CCTTTGCTGATAGTCTCTGCCCAT | TTACCACCAGTCCATCGCCATT |
| sXBP1 | AAGAACACGCTTGGGAATGG | CTGCACCTGCTGCGGAC |
| Dnajb11 | GTACCTCATCGGGACTGTGAT | CAGAACCTCATAAGCAGCACC |
| Edem1 | CGGCTATGACAACTACATGG | GTTCAGATTGGACTCTC |
| Dnajc3 | GGCGCTGAGTGTGGAGTAAAT | GCGTGAAACTGTGATAAGGCG |
| Herpud1 | GCAGTTGGAGTGTGAGTCG | TCTGTGGATTCAGCACCCTTT |
| β-actin | CAGGCATTGCTGACAGGATG | TGCTGATCCACATCTGCTGG |

**Table S2. List of antibodies used for Western blot (WB) and Immunofluorescence (IF).**

| Antibody | Source and Catalog no. | Dilution | Analysis |
|---|---|---|---|
| Polyclonal TWEAK Receptor/Fn14 | Cell Signaling Technology # 4403 | 1:1000 | WB |
| Monoclonal rabbit-anti-GAPDH | Cell Signaling Technology # 2118 | 1:1000 | WB |
| Monoclonal rabbit-anti-phospho-PERK | Cell Signaling Technology Cat# 3179L | 1:500 | WB |
| Monoclonal rabbit-anti-PERK | Cell Signaling Technology Cat# 3192 | 1:500 | WB |
| Monoclonal rabbit-anti-phospho-eIF2α | Cell Signaling Technology # 3398 | 1:500 | WB |
| Monoclonal rabbit-anti-total-eIF2α | Cell Signaling Technology # 5324 | 1:500 | WB |
| Monoclonal rabbit-anti-ATF-4 | Cell Signaling Technology # 11815 | 1:500 | WB |
| Monoclonal rabbit-anti-XBP1s (E9V3E) | Cell Signaling Technology # 40435 | 1:500 | WB |
| Monoclonal mouse-anti-Puromycin | Millipore, MABE343 | 1:500 | WB |
| Monoclonal mouse-anti-MyHC | DSHB #MF 20 | 1:200 | IF |
| Polyclonal rabbit-anti-Laminin | Sigma, L9393 | 1:1000 | IF |
| Polyclonal goat-anti-mouse IgG Alexa Fluor 568 | Invitrogen # A-11004 | 1:1000 | IF |
